# Axon guidance deficits in a human sensory neuron model of Fabry disease

**DOI:** 10.1101/2025.09.01.673441

**Authors:** Christoph Erbacher, Aneeta Andrews, Till Sauerwein, Maximilian Breyer, Panagiota Arampatzi, Maximilian Koch, Stephanie Lamer, Tom Gräfenhan, Andreas Schlosser, Nurcan Üçeyler

**Affiliations:** Department of Neurology, University Hospital Würzburg, Josef-Schneider-Str. 11, 97080 Würzburg; ZB MED – Information Centre for Life Sciences, Cologne, Germany; Core Unit Systems Medicine, University of Würzburg, 97080 Würzburg, Germany; Institute of Clinical Neurobiology, University Hospital Würzburg, Würzburg, Germany; Rudolf Virchow Center, Center for Integrative and Translational Bioimaging, University of Würzburg, Würzburg, Germany

**Keywords:** Fabry disease, induced pluripotent stem cells, disease model, sensory-like neuron, axon guidance

## Abstract

Fabry disease (FD) is a rare genetic galactosidase alpha (*GLA*) gene associated lysosomal disorder caused by alpha-galactosidase A (AGAL) deficiency, leading to sphingolipid (globotriaosylceramide, Gb3) accumulation in multiple tissues. Burning pain due to small fiber neuropathy is an early symptom with great impact on health-related quality of life. The pathophysiological role of Gb3 accumulations in sensory neurons of the dorsal root ganglia is incompletely understood. We have differentiated induced pluripotent stem cells of an isogenic *GLA* knockout line (p.S364del, hemizygous) and its healthy control into sensory neurons to model FD *in vitro*. We have compared both lines on transcriptional and proteomic level and investigated the effects of AGAL enzyme supplementation. FD sensory neurons showed dysregulation of disease-related pathways, including axon guidance at both RNA and protein level and microfluidic assays revealed shorter neurite length. While AGAL did not restore the transcriptomic state, it reduced Gb3 accumulation and lowered protein ephrin 5A and glycoprotein M6A level. These findings highlight axon guidance alterations in an isogenic human FD sensory model, with potential implications for early central and peripheral innervation in small fiber neuropathy.

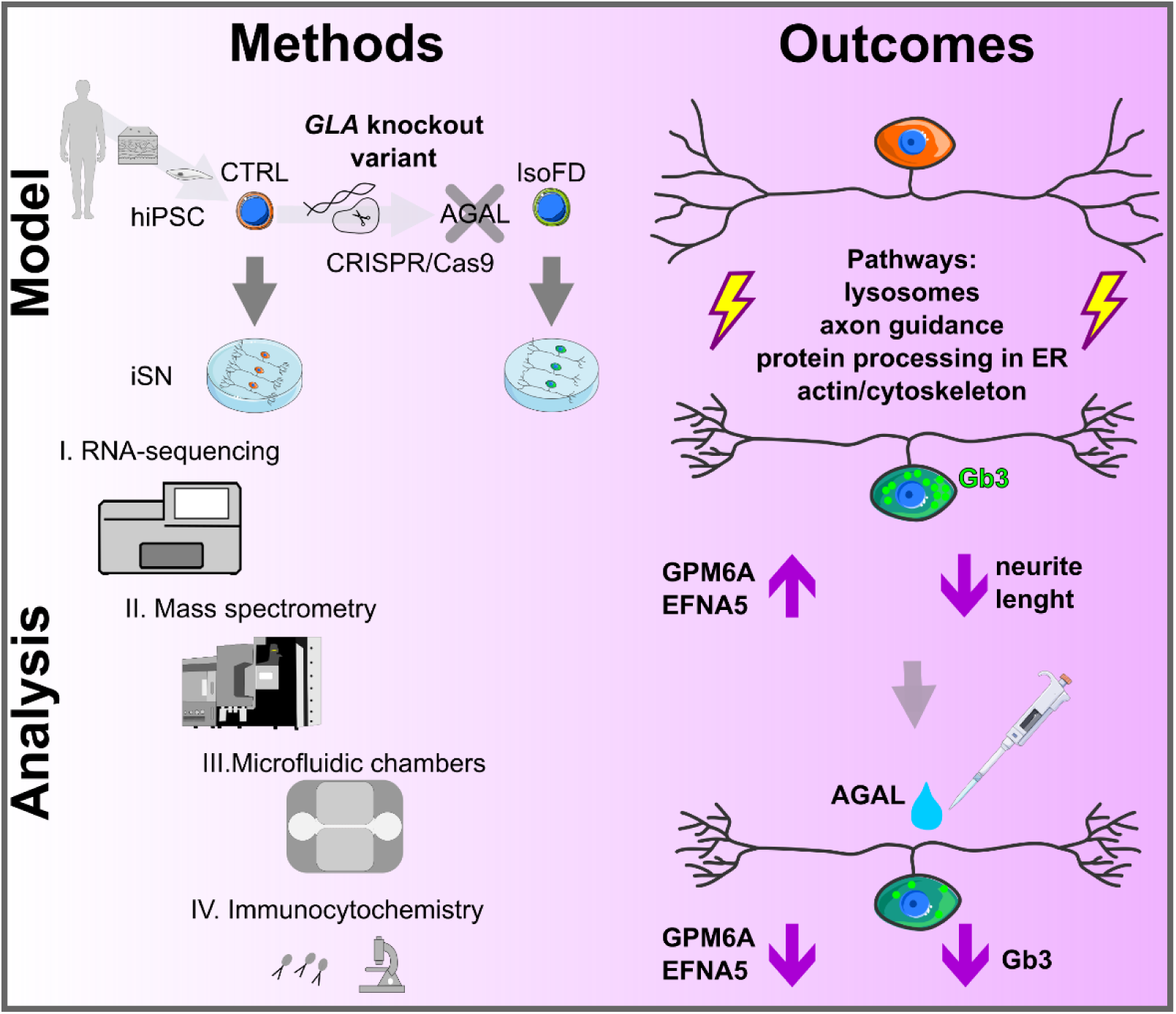

## 1. Introduction

Fabry disease (FD) is a rare X-linked genetic disorder arising from pathogenic variants in the galactosidase alpha (*GLA*) gene [1]. These variants lead to the deficient or absent activity of the lysosomal enzyme alpha-galactosidase A (AGAL), which is responsible for hydrolyzing terminal alpha-galactosyl moieties from glycolipids, particularly globotriaosylceramide (Gb3) [2]. As a result, Gb3 and other sphingolipids accumulate in lysosomes, causing a multisystem disorder with severe cardiac and renal complications [3, 4]. Episodic acral burning pain is one of the earliest FD symptoms based on small fiber neuropathy (SFN) [5]. Other manifestations of SFN in FD include loss of cold sensation and skin denervation [6, 7]. Current treatments aim to reduce FD symptoms by enzyme replacement, restoration of residual enzyme function via chaperone, or substrate reduction [8].

Whilst all these approaches can slow disease progression when started early, they are not curative [7, 9, 10].

The pathophysiology of SFN in FD is incompletely understood. Animal models and human studies show that Gb3 accumulates in various cell types, including sensory neurons of the dorsal root ganglia (DRG) [11]. In animals, these accumulations were linked to altered ion channel function, (neuro-)inflammation, and hypoxic environment at DRG level [12–15], yet discrepancies across species hamper translation of findings towards patients [16, 17]. Access to native human sensory neurons or DRG is highly restricted, limiting direct insights on SFN in FD patients.

Induced pluripotent stem cell (iPSC) -based sensory-like neuron (iSN) models [18] can bridge this gap and characterized iSN from FD cell lines revealed electrophysiological alterations, increased calcium flux, and neurite thinning [19]. Still, the cellular mechanisms that drive these pathological changes are unclear. Here, we analyzed iSN from a FD iPSC line carrying the pathogenic *GLA* variant versus its isogenic healthy control on a molecular basis. We applied transcriptomic and proteomic analyses and uncover alterations in axon guidance pathways, validated by immunolabeling and microfluidic neurite outgrowth experiments.

## 2. Materials and methods

### 2.1. Cell culture

Established iPSC of a male healthy control line (CTRL) and its isogenic *GLA* knockout line (IsoFD; p.S364del, hemizygous; introduced via CRISPR/CAS9), were previously generated after ethical approval (Ethics Committee of the University of Würzburg Medical Faculty; #135/15) [19]. iPSC were cultured at 5% CO_2_ and 37°C in 2 ml MACs medium with supplement (Miltenyi Biotec, Bergisch Gladbach, Germany) and 100 U/ml pen/strep (Thermo Fisher Scientific, Waltham, MA, USA) with daily medium change . We differentiated iPSC into iSN according to an established protocol [19, 20] (see Supplementary materials) and regularly screened iPSC for mycoplasma contamination (Fig. S1). Neurons were cultivated for 6 weeks with either maturation medium or maturation medium with 1.32 μg/ml α-galactosidase A; (AGAL; Sanofi Genzyme, Cambridge, MA, USA) added every 14 days.

### 2.2. Transcriptomics

Maturation medium was removed from one 6-well per cell line and condition, and iSN were mechanically detached via up- and down pipetting with warm DMEM/F12 medium (Thermo Fisher Scientific, Waltham, MA, USA) via a 10-ml pipette.

iSN were transferred into a 15 ml falcon and pelletized at 300 x g for 3 min. Medium was removed and cells lysed in 700 µl Qiazol (Qiagen, Hilden, Germany) with up and down pipetting and brief vortexing. In addition, remaining adhesive cells in the 6-well were lysed in Qiazol. Afterwards, the miRNeasy Mini Kit (Qiagen, Hilden, Germany) was used to extract total RNA. Samples were stored at -80°C until further processing. RNA from four independent differentiations for each condition (CTRL, CTRL-AGAL, IsoFD, IsoFD-AGAL) was analyzed, except for CTRL-AGAL with four samples from three differentiations. RNA was quality-controlled on a 5200 Fragment Analyzer with the DNF-471-33 - SS Total RNA 15nt kit (Agilent Technologies, Santa Clara, CA).

The RIN for all samples was >7. Details regarding the library preparation part and sequencing are included in the supplementary material. Separate DESeq2 analyses were conducted to achieve different data resolutions, once considering sample assignment to conditions CTRL, CTRL-AGAL, IsoFD, and IsoFD-AGAL (see Supplementary materials). The RNA-Seq data presented in this work has been deposited to the European genome-phenome archive (EGA; EGA ID: will be publicly available as of the date of publication). DESeq2 results are shown in Supplementary Tables S1-S4. Differential expressed (DE) genes were assumed at an adjusted P- value (padj) <0.05 after Benjamini–Hochberg correction.

### 2.3. Proteomics

Maturation medium from two 6-wells per cell line and condition was removed and iSN were mechanically detached via up and down pipetting with ice cold PBS (Thermo Fisher Scientific, Waltham, MA, USA). Neurons from both wells were pooled and pelletized at 300 x g for 3 min.

Medium was removed and cells lysed in RIPA buffer (Thermo Fisher Scientific, Waltham, MA, USA) with 1x protease inhibitor (cOmplete, EDTA-free; Roche, Basel, Switzerland) under vigorous pipetting for 1 min followed by incubation on ice for 4 min. Afterwards, samples were flash frozen in liquid nitrogen and stored at -80°C until further processing. Three independent differentiations for each condition (CTRL, IsoFD, IsoFD-AGAL) were used. Before gel electrophoresis, protein samples were thawed and centrifuged at 14.000 x rcf for 15 min at 4° and the supernatant transferred to a new Eppendorf tube. A bicinchoninic acid assay (BCA) Protein Quantitation Kit (Interchim, San Diego, CA, USA) was used to estimate protein concentration via a NanoDrop Onec spectrophotometer Thermo Fisher Scientific, Waltham, MA, USA) according to the manufacturer’s protocol. Samples were subjected to mass spectrometry according to an established protocol (see Supplementary materials). MaxQuant was used to analyze Raw MS data files[21]. To identify deregulated proteins, mean log2-transformed protein ratios of FD versus CTRL samples were derived from the three replicate experiments. In addition, the R package limma [22] was used to calculate Benjamini-Hochberg adjusted p-values and FDR. Proteins with a padj <0.05 were considered deregulated. All proteins with uncorrected p <0.05 were used for exploration of enriched pathways.

### 2.4. Downstream bioinformatics analysis

For transcriptomic pathway analysis, DESeq2 datasets were applied for gene set enrichment analysis (GSEA) performed with clusterProfiler version 3.12.0 [23] using the pathways from the Kyoto Encyclopedia of Genes and Genomes. A false discovery rate q-value <0.01 was chosen to identify pathway expression differences. Only protein coding genes were considered because these are best annotated for pathway analysis.

Metascape [24] was used for over-representation analysis (ORA) to derive enriched terms and cell type signatures for CTRL iSN versus CTRL feeder-like cells (iF) by applying the genes with padj <0.01 and log2f change >4 (enriched in iF) and log2f change < -4 (enriched in iSN). Deconvolution of iSN bulk-RNA was performed via BayesPrism [25], using raw read counts per gene from each iSN line and condition and a published single cell sequencing dataset from human DRG as reference profile [26]. The bioinformatical workflow of the BaysPrism analysis is publicly available on Zenodo (https://doi.org/10.5281/zenodo.15045467)

For proteomics, metascape ORA was used, applying all proteins with p <0.05 from limma for each condition. Principal component analysis (PCA) was computed via ClustVis [27]. TPM (RNA transcripts) and log10LFQ (protein) values for corresponding conditions and differentiations for matching transcript-protein pairs were correlated via Spearman rank test in R [28].

### 2.5. Immunocytochemistry and image analysis

We validated key transcriptomic and proteomic targets linked to axon guidance and neurite outgrowth via immunocytochemistry (ICC). Six-week old neurons in 8-well chambers (C8-1.5H-N; Cellvis, Mountain View, CA, USA) were fixed with 4% PFA in PBS for 15 min at RT, followed by 5 min washing with PBS twice. Primary antibodies against ephrin 5A (EFNA5; #AF3743, Lot: BQB042405A, R&D Systems, McKinley Place NE, MN, USA) and neuronal membrane glycoprotein M6-A (GPM6A; #D055-3, Lot: 027, MBL Life Science, Tokyo, Japan), were applied 1:200 on iSN over night at 4°C in an antibody solution with 0.1% (v/v) saponin and 10% (v/v) fetal calf serum (FCS) in PBS.

After washing with PBS twice, secondary antibodies #712-545-150 and #705-165- 147, Dianova, Hamburg, Germany) and anti-peripherin (PRPH; # sc-377093, Lot: F1323 Santa Cruz Biotechnology, Dallas, TX, USA) conjugated with Alexa Fluor 647 were applied 1:200 for 30 min in PBS at RT. Alternatively, iSN were first incubated without primary antibodies, followed by anti-PRPH (# sc-377093, Lot: D1818, Santa Cruz Biotechnology, Dallas, TX, USA) conjugated with Alexa Fluor 488 at 1:200 and shiga toxin conjugated to Atto643 [19] at 1:500 for 30 min in PBS at RT. Washing steps were applied as for first antibodies. Subsequently, neurons were kept in PBS and stored at 4°C until imaging. Neurons were imaged with a ThunderImager equipped with a K5 monochrome camera using a 20x magnification objective via LasX software (all: Leica Microsystems, Wetzlar, Germany). Three differentiations per group, each in duplicates were imaged with six non-overlapping region of interest (ROIs) per well. Imaging was performed blinded for the differentiation and iSN group and z-focus controlled only by the PRPH channel. Fiji pipelines, leveraging the PRPH signal as predetermined iSN ROI (to exclude iF) were used. For GPM6A and EFNA5, the PRPH area was skeletonized prior to measuring intensity levels of targets to exclude bias by differently sized soma clusters (see Fiji pipelines in Supplementary materials).

### 2.6. Microfluidic culture of iSN and outgrowth stimulation

Microfluidic chambers (Xona Microfluidics, SND 150, Research Triangle Park NC, USA) were coated in two steps: first with 1 mg/ml poly-L-lysine (Sigma Aldrich, St. Louis, MO, USA) in ddH2O over night at RT, followed by 2.5 μg/ml laminin in ddH2O (Thermo Fisher Scientific, Waltham, MA, USA) for at least 3.5 h at RT.

Briefly, iSN were detached after differentiation and were resuspended after centrifugation in 1 ml maturation medium including 10 μM Y27632 (Miltenyi Biotec, Bergisch Gladbach, Germany). To enrich for iSN, cells were plated on bMg-coated 6- wells with 2 ml of maturation medium and settled in an incubator for 20 min.

Subsequently, the remaining floating cell fraction was transferred into a 15-ml falcon tube and centrifuged at 330 x g for 3 min. The final pellet was resuspended in 60 µl fresh maturation medium with 10 μM Y27632 (Miltenyi Biotech, Bergisch Gladbach, Germany). The central compartment of each chamber was filled with 10 μl of cell suspension. Adjacent compartments contained maturation medium with 40 ng/ml human recombinant beta-nerve growth factor (bNGF; PeproTech, Rocky Hill, NJ, USA). Neurons were cultured for 7 days with medium change every third day and subsequently fixed with 4% paraformaldehyde (PFA) for 15 min at RT. Subsequently, iSN were incubated over night with 1:200 anti-PRPH (# sc-377093, Lot: D1818, Santa Cruz Biotechnology, Dallas, TX, USA) conjugated with Alexa Fluor 488 in PBS with 0.1% (v/v) saponin and 10% (v/v).

Microfluidics were washed twice with PBS and imaged as indicated above, with a 10x objective (Leica Microsystems, Wetzlar, Germany). CTRL and IsoFD iSN from four differentiations were analyzed each. Total neurite length per compartment was quantified in a blinded manner via manual tracking with NeuronJ [29]. Only neurites crossing through the microfluidic barrier were counted and only compartments with outgrowth analyzed.

### 2.7. Statistical analysis and graphs

Imaging data were analyzed with SPSS software version 29 (IBM, Armonk, NY, USA). Normal distribution was assessed using the Kolmogorov–Smirnov and the Shapiro–Wilk test. The two-tailed Mann-Whitney U test was used for comparing neurite outgrowth between two independent groups. Pairwise one-tailed Mann-Whitney U tests with Holm-Sidak post-correction were applied for multiple comparisons between immunolabeled iSN groups. Data were visualized as box-and-whisker plots using GraphPad PRISM version 8 (GraphPad Software, Inc., La Jolla, CA, USA). Transcriptome related volcano plots were plotted in R (version 4.4.1) with ggplot2 and dplyr [28]. Figures were compiled in Inkscape.

## 3. Results

### 3.1. Molecular characterization of sensory-like neurons

iSN corresponded to sensory neurons as evidenced by high gene marker expression compared to the more adhesive non-neuronal ‘feeder-like’ (iF) cells (Fig. 1A).

**Figure 1.**
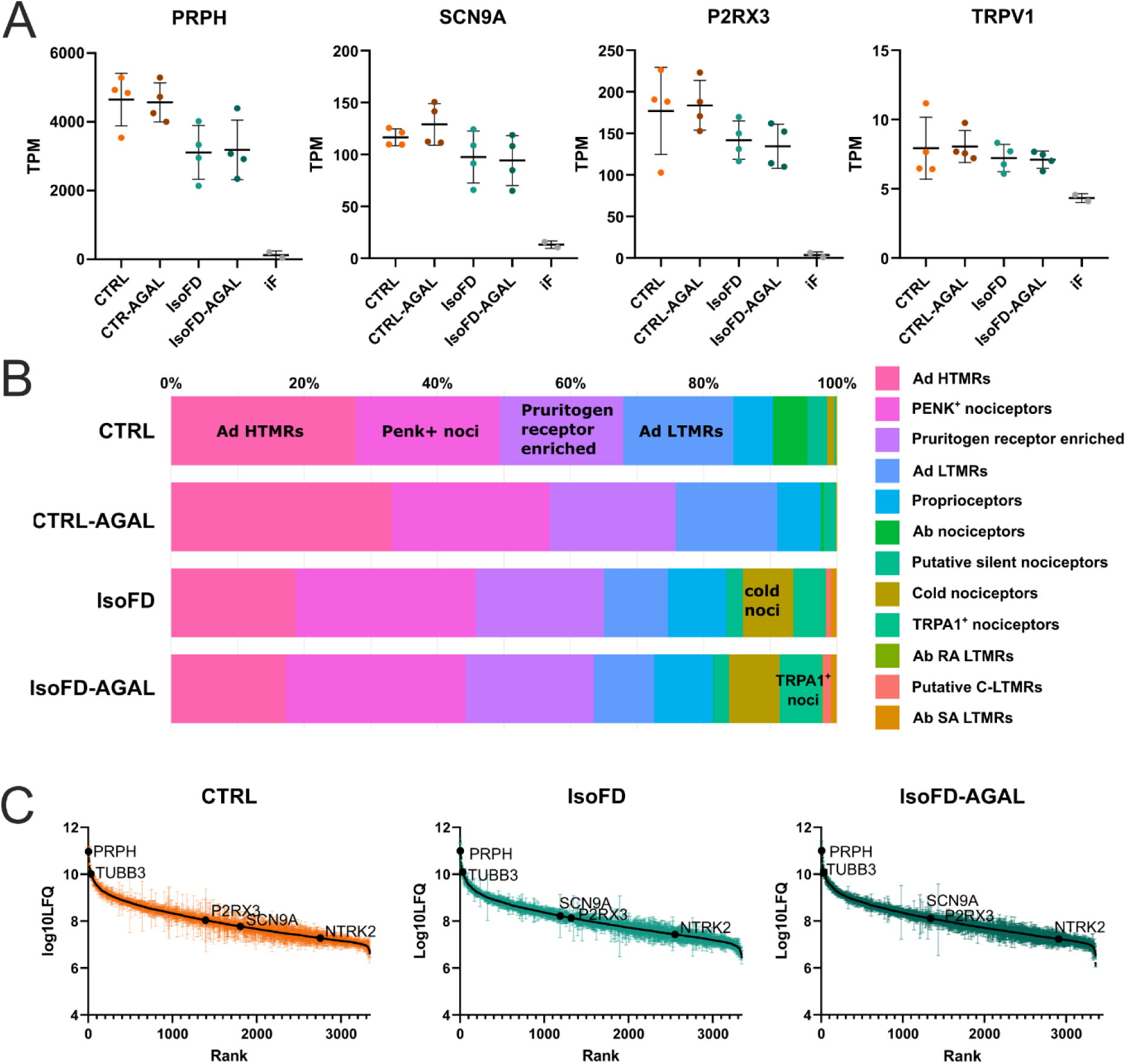
Molecular characterization of iSN. A) RNA expression levels of peripheral neuronal marker *PRPH*, sensory neuron-specific marker sodium voltage-gated channel alpha subunit 9 (*SCN9A*), sensory neuron subtype-specific purinergic receptor P2X 3 (*P2RX3*), and transient receptor potential cation channel subfamily V member 1 (*TRPV1*) were consistently expressed across differentiations, cell lines, and treatment conditions (n=4 differentiations per line and treatment). In contrast, these markers showed low expression in CTRL iF (n=2). B) Deconvoluted cell-type proportions of iSN groups (mean percentage from each n=4 differentiations). C) Ranked display of detected proteins showed comparable levels of detected sensory neuron markers.

Deconvolution via a human DRG spatial sequencing dataset [26] revealed highest similarity of iSN to Ad HTMRS, PENK+ nociceptors, pruritogen receptor enriched, and Ad LTMRs DRG neuron subtypes, with minimal proportional differences between CTRL and IsoFD iSN, irrespective of AGAL treatment (Fig. 1B). Mechanically detached iSN showed distinct transcriptomic divergence from the remaining adhesive non-neuronal iF (Fig. S2). Highly DE transcripts (log2f < -4, padj <0.01) distinguishing iSN from iF aligned with DRG gene expression and were enriched in neuronal pathways (Fig. S2). Overall, protein expression showed consistent expression of characteristic sensory neuron markers in IsoFD and CTRL lines, not affected by AGAL treatment (Fig 1C).

### 3.2. FD iSN show transcriptomic alterations

When comparing IsoFD iSN with CTRL iSN, 297 transcripts were differentially expressed, mainly protein coding (66.3%), but interestingly also many lncRNAs (23.2%) transcripts (Fig. 2A, Supplementary Table S1). AGAL showed only a mild effect on the IsoFD transcriptome, with 10 upregulated ribosomal RNAs compared to untreated IsoFD (Fig. S3, Supplementary Table S2). Thus, comparing IsoFD-AGAL versus CTRL revealed 260 DE transcripts (Fig. S3, Supplementary Table S3).

**Figure 2.**
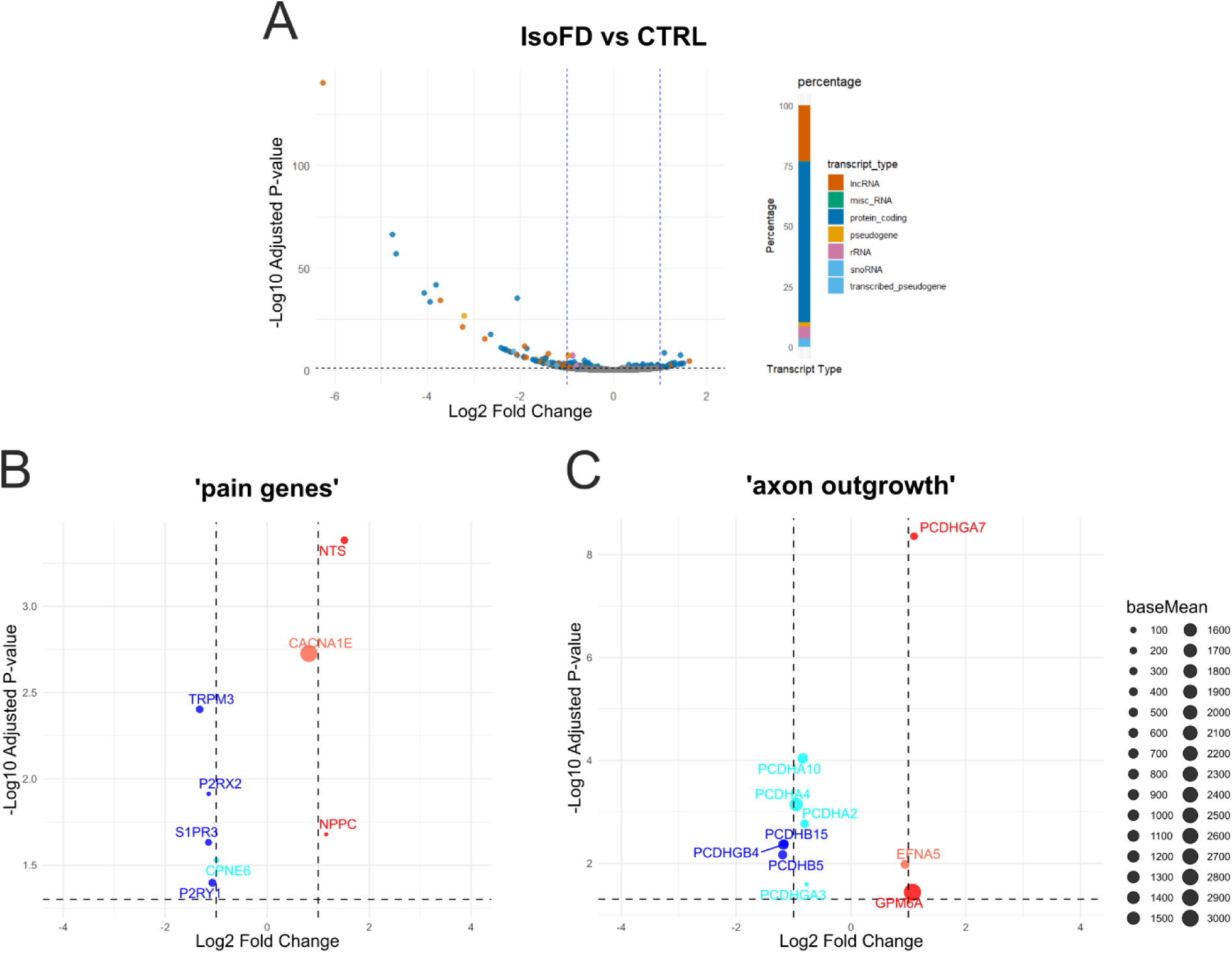
Transcriptional deregulation in FD iSN. A) Volcano plot with DE of transcripts from IsoFD versus CTRL. B) DE protein coding genes in IsoFD versus CTRL iSN related to pain and C) related to axon guidance and outgrowth with their baseMean expression level indicated by dot size. Log2 fold change < -1 = blue, Log2 fold change > -1 < 0 = cyan, Log2foldchange > 0 < 1 = orange, Log2fold change > 1 = red. Data from n=4 differentiations each with padj <0.05.

Finally, AGAL showed no impact on healthy CTRL iSN with only one DE transcript (Fig. S3, Supplementary Table S4). Transcriptomic differences between IsoFD and CTRL iSN prompted further exploration of IsoFD-related changes. From a published list of 486 known pain-related genes [30] we found 10 DE genes in IsoFD (Fig. 2B). We also identified Ephrin A5 (*EFNA5*), glycoprotein M6A (*GPM6A*), and eight protocaderins (Fig. 2C) as DE genes involved in axon guidance and outgrowth [31–33].

### 3.3. Gene set enrichment identifies FD- and neuron-specific pathways

To more systematically characterize FD-related changes in iSN, we applied GSEA which revealed 24 enriched KEGG pathways, mostly showing predominant gene upregulation (Fig. 3A). Notably, the top ten included pathways relevant to FD, such as lysosome, protein processing in endoplasmatic reticulum, PI3K-Akt signaling, and, critically for sensory neurons, axon guidance (Fig. 3B, Supplementary Table S5).

**Figure 3.**
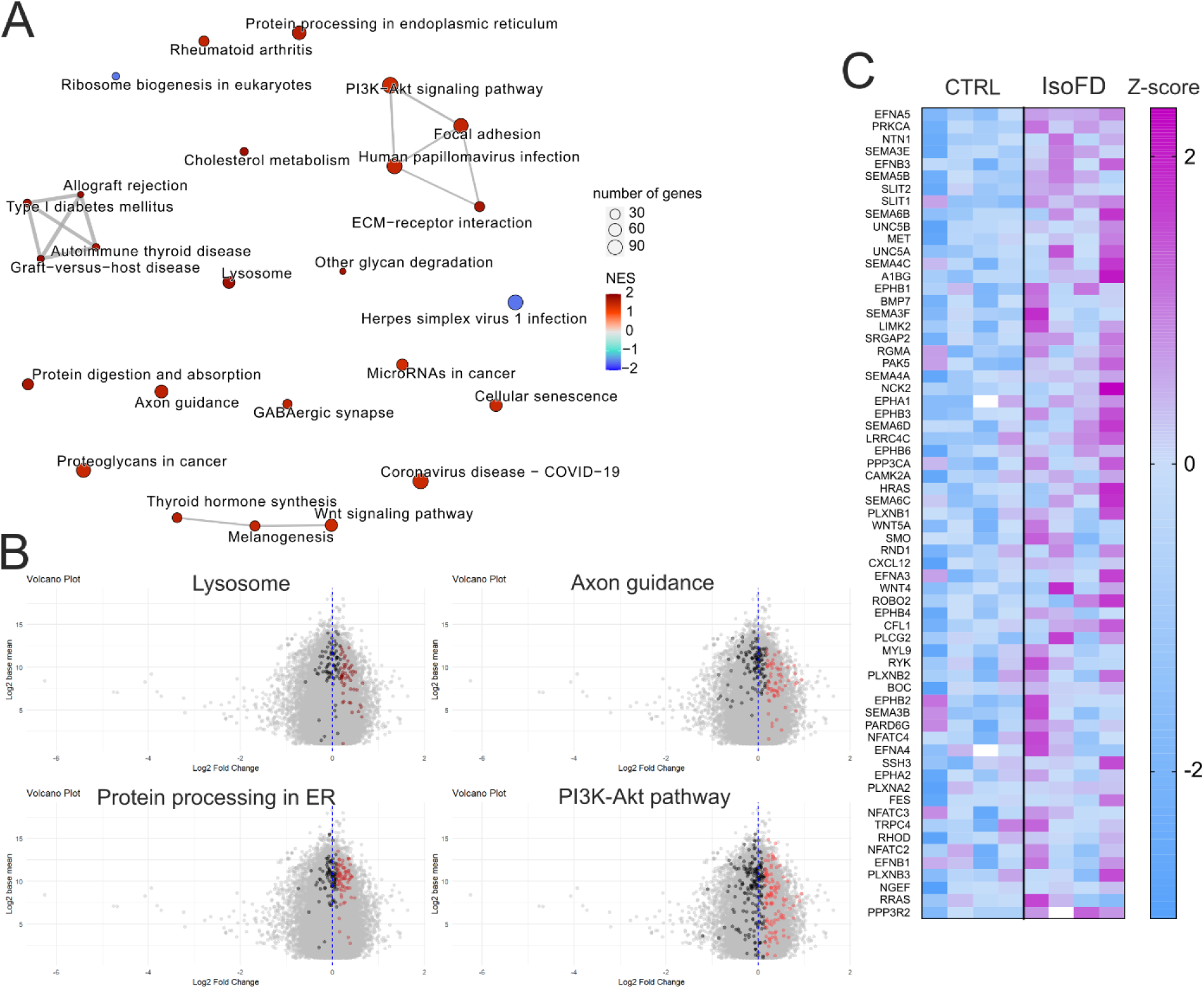
Gene set enrichment analysis of IsoFD versus CTRL iSN transcriptome. A) Enriched pathways and their connections shown with positive enrichment (red) and negative enrichment (blue). All pathways with q<0.05 shown. B) Representative pathways within the top 10 displaying unrelated genes (grey), core enriched genes (colored), and not enriched genes of the respective pathway (black). C) Changes in expression of axon guidance pathway associated genes. Data from comparison of n=4 differentiations each. To calculate z-scores, normalized means were log10- transformed and the following formula was applied: z-score = (log10norm.count of sample - mean log10normcount of CTRL) / SD of log10norm.count CTRL.

### 3.4. IsoFD iSN proteome reflects cytoskeleton and axon guidance adaptions

We sought to corroborate our transcriptomic results at the protein level via mass spectrometry, yet PCA did not reveal a clear group separation (Fig. 4A). Differential protein expression analysis showed no dysregulation between IsoFD and CTRL iSN after post-correction (Fig. 4B, Supplementary Table S6). Since only a fraction of all protein coding transcripts was measured at protein level and only 15/197 (7.6%) of DE coding genes were identified (Fig. 4C), we did not apply GSEA. Instead, we performed ORA using all uncorrected p<0.05 proteins via metascape. Within the enriched terms, ‘actin-filament based processes’ and ‘axon guidance’ emerged (Fig. 4D, see Supplementary Table S7), suggesting conserved alterations of neurite outgrowth in IsoFD at protein level. Side-by-side comparison of the protein lists of IsoFD-AGAL and IsoFD, revealed lesser enrichment for these terms in IsoFD-AGAL and ‘translation’ was only enriched as a pathway in these iSN (Fig. 4D). Still, Differential protein expression analysis between IsoFD and AGAL-treated IsoFD revealed only one differentially expressed protein after post-correction, namely AGAL (gene identifier *GLA*), indicating the effective uptake of supplemented enzyme in culture (Supplementary Table S8). Expression levels of transcripts and detected proteins of iSN from the same differentiations correlated moderately and this correlation strength was not altered in IsoFD (Supplementary Fig. S4).

**Figure 4.**
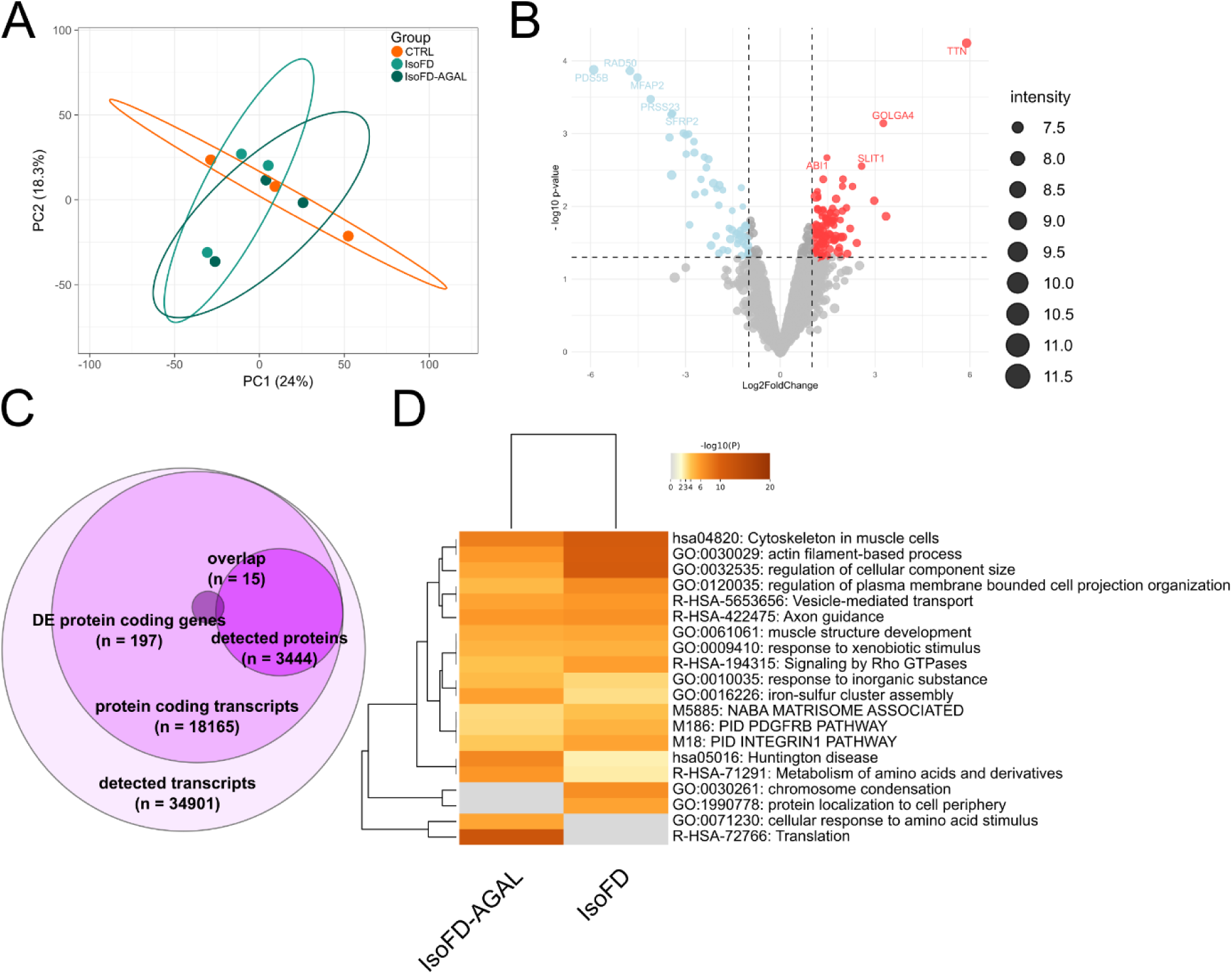
Indicators of axon guidance and actin-related alterations in FD iSN on proteome level. A) PCA showed low group separation. B) Differences in protein abundance between IsoFD and CTRL iSN at p<0.05. C) Limited overlap between detected RNA transcripts and corresponding proteins. D) Actin filament-based process and axon guidance were altered in IsoFD according to ORA analysis, with alterations partially mitigated in AGAL supplemented IsoFD (cutoff for pathways at q<0.05). Data from 3 differentiations each.

### 3.5. IsoFD iSN mean neurite length is reduced

To test whether shifts in axon guidance pathways affect neurite outgrowth in IsoFD, we cultured iSN in microfluidic chambers (Fig. 5A). After one week of bNGF stimulated outgrowth total neurite length in the outer compartments remained unchanged (Fig. 5B-C). However, IsoFD iSN showed shorter mean neurite length, suggesting increased arborization but reduced elongation.

**Figure 5.**
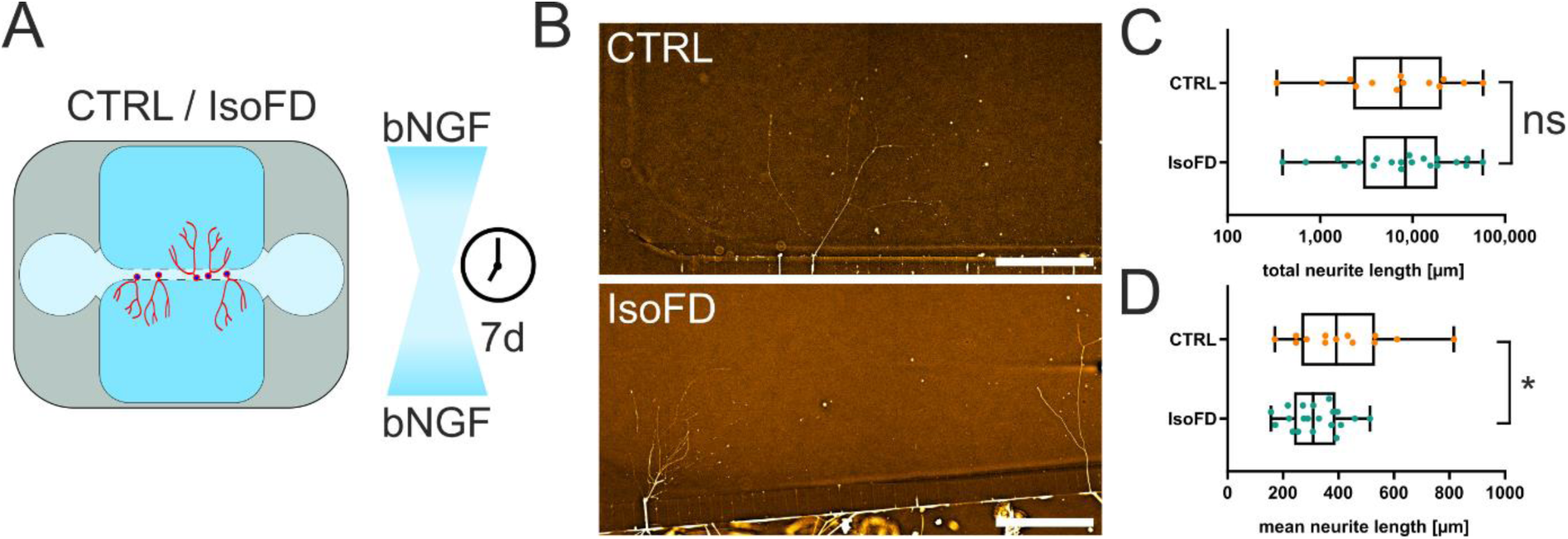
Microfluidic outgrowth assay indicates altered neurite outgrowth of IsoFD iSN. A) Overview of microfluidic chamber culturing for seven days with bNGF gradient. B) Examples of neurite outgrowth visualized by PRPH labeling. C) No difference in total neurite length per compartment between IsoFD and CTRL iSN (p = 0.813). D) Reduced mean neurite length per compartment in IsoFD compared to CTRL iSN (p =0.041). Comparison of compartments (n_CTRL_ = 13, n_IsoFD_ = 20) derived from n=4 differentiations each, using unpaired two-tailed Mann-Whitney U test (C) or unpaired t-test (D). *p < 0.05, ns = not significant. Each dot represents one chamber compartment.

### 3.6. Axon guidance proteins GPM6A and EFNA5 are elevated in IsoFD iSN and reduced by AGAL supplementation

To validate our transcriptomic and proteomic findings on axon guidance and neurite outgrowth, we assessed key proteins via ICC. While no suitable antibodies for protocaderins were found, GPM6A and EFNA5 antibodies were previously validated [34, 35]. Gb3 accumulation was quantified via shiga toxin labeling [19] to confirm activity and its effect on candidate proteins. IsoFD showed increased GPM6A and EFNA5 expression (Fig. 6 A-B), which AGAL treatment reduced to CTRL-like levels. AGAL activity was evident by a strong Gb3 reduction in IsoFD-AGAL, though not fully restored to CTRL levels (Fig. 6C).

**Figure 6.**
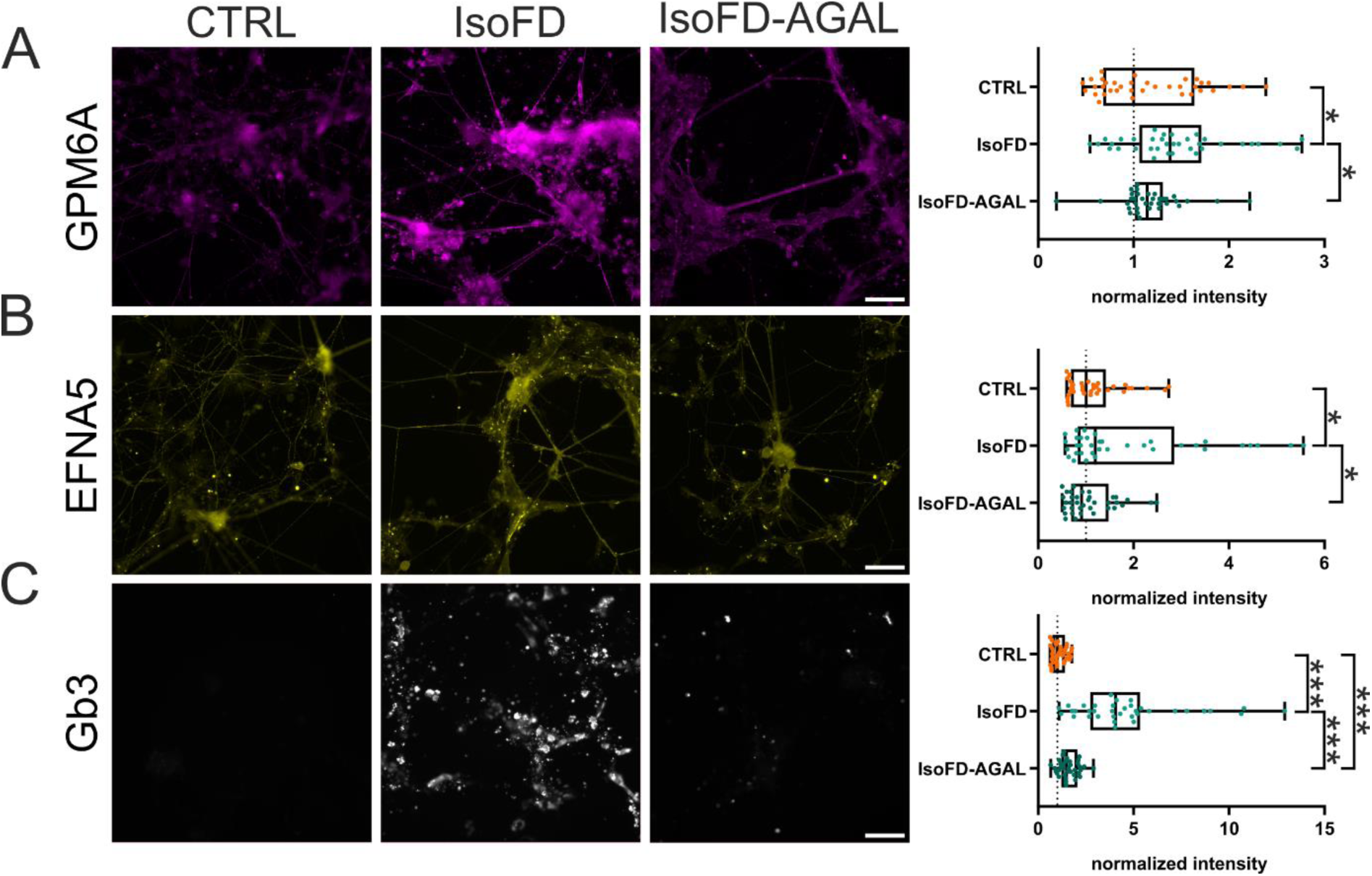
Validation of axon guidance protein deregulation and alleviating effect of AGAL. A) Higher expression of GPM6A in IsoFD iSN versus CTRL iSN (p = 0.034) and versus IsoFD-AGAL iSN (p = 0.023). B) Higher EFNA5 expression in IsoFD iSN versus CTRL iSN (p = 0.046) and IsoFD-AGAL iSN (0.018). C) IsoFD showed higher Gb3 signal compared to CTRL. AGAL treatment decreased Gb3 signal in IsoFD- AGAL iSN compared to IsoFD iSN, yet did not completely restore levels compared to CTRL iSN (IsoFD versus CTRL p < 0.001, IsoFD-AGAL versus IsoFD p < 0.001, IsoFD-AGAL versus CTRL p < 0.001). Scale bar: 100 µm. Comparison between each three differentiations with each two wells and six images per well (nCTRL, nIsoFD, nIsoFD-AGAL = 36 images each). Each dot represents one image. One-tailed Mann-Whitney-U test with Holm-Sidak correction. *p < 0.05, **p < 0.01, ***p < 0.001.

## 4. Discussion

We investigated molecular mechanisms of small fiber pathology in FD using a human iSN model. Our findings revealed multilevel evidence for axon guidance deficits in the isogenic FD model cell line, both at transcriptional and proteomic level and via a functional outgrowth assay.

We estimated cell type proportions using bulk RNA-seq deconvolution, leveraging a human DRG subtype matrix. Our iSN were comparable to iPSC-derived sensory-like neurons with another differentiation approach [36]. Although no shift in the major DRG neuron subtype similarities was present in CTRL and IsoFD iSN with or without AGAL treatment, interestingly the percentage of cold nociceptors and transient receptor potential cation channel subfamily A member 1 (TRPA1)-positive nociceptors was increased in IsoFD.

Reduced epidermal innervation is one hallmark of SFN and displays a profound and early sign in FD patients [37]. Axon guidance molecules and their receptors and protocaderins are crucial in innervation and orchestrate the development of embryonic DRG and directing their targeting to end organs [38, 39], while their post-developmental role is less investigated. On RNA level, several protocadherins, EFNA5 and GPM6A were deregulated in IsoFD iSN.

GPM6A regulates neurite outgrowth, filopodia/dendritic spine formation, and synapse formation in the central nervous system, and is associated with lipid rafts linked to sphingolipids [40, 41]. It was recently implicated in sensory neuron neurite outgrowth [34]. We observed increased GPM6A expression at both RNA and protein levels, aligning with a study reporting membrane-specific upregulation in another FD iSN line [42]. EFNA5, known for repulsive interactions, is linked to growth cone collapse and neurite redirection in sensory neurons [43].

iSN may represent less mature sensory neurons compared to postnatal animal models or organ donor DRG neurons [37, 44], raising the possibility that FD could impact early development, not just age-related nerve fiber loss. Small fiber-related symptoms often present in pediatric patients [45], and reduced skin innervation was observed in 2-3 month old FD model mice [12, 46], but data at the embryonal stage are lacking. FD may affect sensory neuron subtype development differentially, with A- beta fibers less impacted than A-delta and C-fibers [47]. However, our bulk RNA sequencing and differences between 3D *in vivo* and 2D *in vitro* models preclude definite conclusions.

The second hallmark of SFN in FD is loss of mainly cold sensation along with evoked and spontaneous acral pain [6, 48]. We found moderate changes in the transcription of several genes linked to pain including *NTS*, *CACNA1E* (both upregulated), *TRPM3*, and *P2RX2* (both downregulated). GSEA showed enrichment of the PI3K- Act pathway, known to play a role in nociception and pain generation [49], which may contribute to altered neuronal excitability in FD [16].

In general, IsoFD iSN did not display a clear “pain signature”. Notably, non-neuronal peripheral cells and immune cells at the skin [50], nerve [51] and DRG levels [52], which influence neuronal activity, were absent in our model. Additionally, synaptic wiring between sensory neurons and dorsal spinal cord neurons, which modulate sensory and nociceptive signal transmission [53], was not present here. Finally, molecular changes in sensory neurons that lead to sensory deficits and pain may evolve over longer periods, potentially undetectable in our 6-week old iSN.

It is crucial to identify the drivers of these changes to uncover potential targets for pharmaceutical intervention. GSEA revealed additional cellular pathways affected in IsoFD iSN, including lysosomes, PI3K/Akt-signaling pathway, and protein processing in the ER. Our CRISPR/Cas9 approach generated a non-functional, but translated AGAL, which may contribute to pathology via unfolded protein response in the ER, as described in other models [54, 55]. However, signs of apoptosis, observed in aged AGAL-deficient mouse sensory neurons [12], were not apparent in our FD iSN model on transcriptomic and proteomic level. We suggest that early deficits in lysosomal and ER function may disrupt nerve fiber outgrowth. Early interventions to enhance lysosomal function and clear non-functional AGAL, in addition to AGAL-based enzyme replacement therapy, could protect DRG neurons and alleviate SFN symptoms in FD.

Biweekly AGAL treatment did not restore transcriptional changes in IsoFD, possibly due to short-term effects or insufficient supply or uptake by iSN. Different cell types exhibit varying capacities to incorporate AGAL [56]. However, our proteomics analysis showed a clear increase in detected AGAL and reduction in cellular Gb3 load after AGAL incubation of iSN, consistent with findings from a previous study [19].

Other factors limiting the curative effect of AGAL could include partial cleavage of Gb3, a short window of enzymatically active AGAL, or the additional pathological impact of non-functionally translated AGAL. However, on protein level, AGAL supplementation reduced GPM6A and EFNA5 levels, suggesting post-transcriptional or translational regulation.

A major limitation of our study is the contrast between >1000 known *GLA* variants [57] and our single FD iPSC line. It is important to note the vast heterogeneity in both genetic and phenotypic backgrounds among FD patients. While cell culture experiments cannot fully address this diversity, we aimed to reduce variability by comparing one healthy control line with one isogenic FD line modeling complete AGAL loss and residual translation. This approach recapitulates a phenotype typical of classical FD caused by missense mutations. Another limitation is the use of a 2D microfluidic outgrowth assay, which oversimplifies complex axon guidance processes and inherits variability in terms of seeding, attachment, and positioning. Further, outgrowth was analyzed after one week of maturation, as prolonged culturing (to a mature 6-week state) was not feasible with the available coating strategies. Finally, during differentiation, varying proportions of iPSC undergo divergent development towards non-neuronal cells [58], here termed iF. Often these confounders are not taken into account during analysis of differentiated neurons. Here, to avoid hybrid transcriptomes and proteomes of iSN and iF, which could obscure molecular iSN profiles, we mechanically detached iSN before lysis. Still, we cannot rule out a small amount of iF attached to neuronal networks during separation.

We present a comprehensive molecular profile of the effects of a *GLA* mutation and subsequent Gb3 accumulation in sensory-like neurons, highlighting human iPSC-based models as a key tool for understanding and addressing the neurological aspects of FD.

## Acknowledgments

We thank the Fabry Center for interdisciplinary Therapy (FAZiT), University Hospital Würzburg, Germany and especially Irina Schumacher for expert technical help. We thank the Core Unit Systems Medicine at the University of Würzburg for excellent technical support and RNA-seq data generation. The Graphical Abstract was partly generated using SMART - Servier Medical ART templates, provided by Servier, licensed under CC BY 4.0.

## Supplementary Figures

**Supplementary Figure S1.**
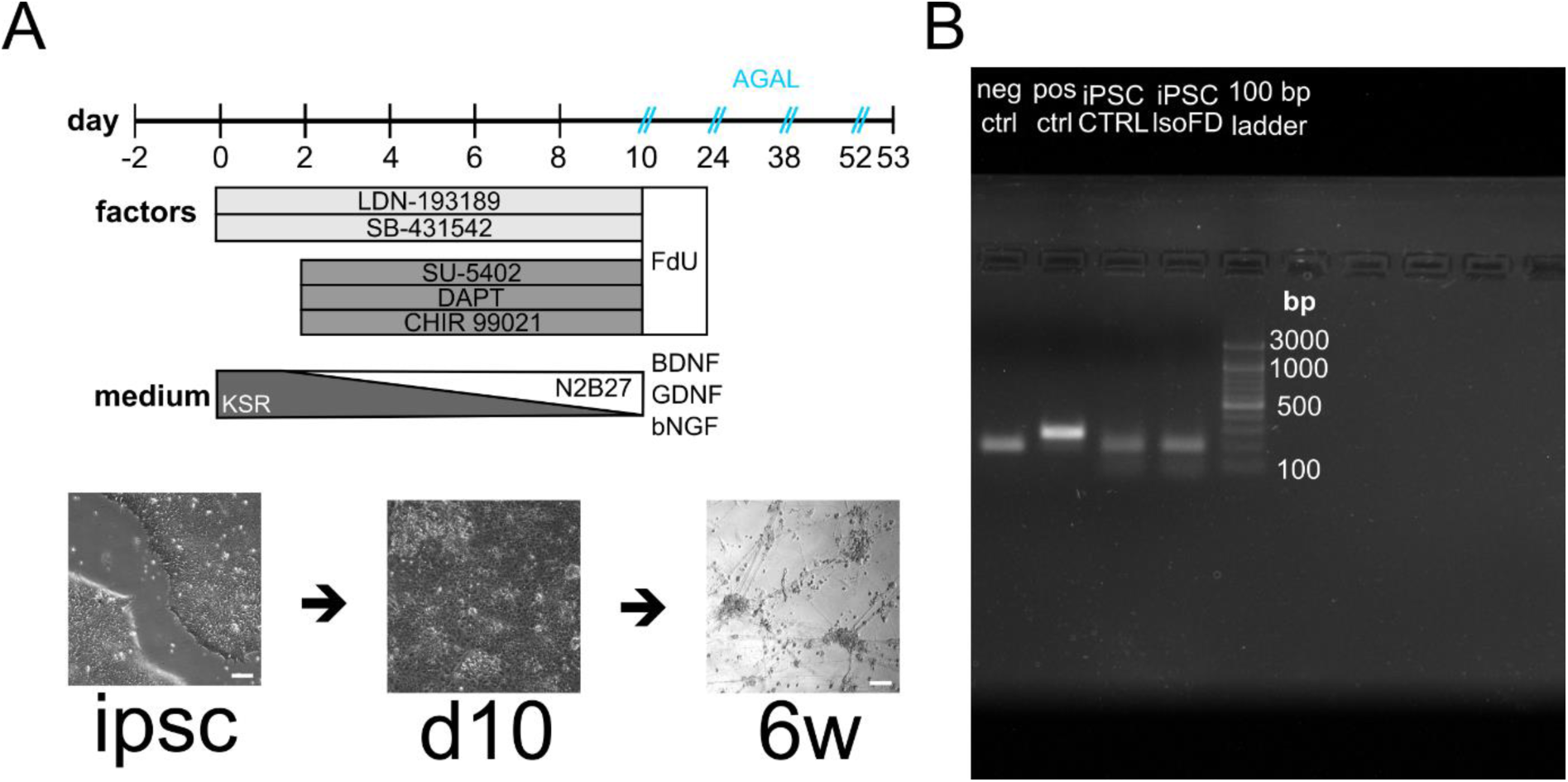
Neuronal differentiation. A) Differentiation protocol with added factors over time and timepoints of AGAL supplementation. B) Screening for mycoplasma DNA via PCR in applied cell lines indicated no contamination.

**Supplementary Figure S2.**
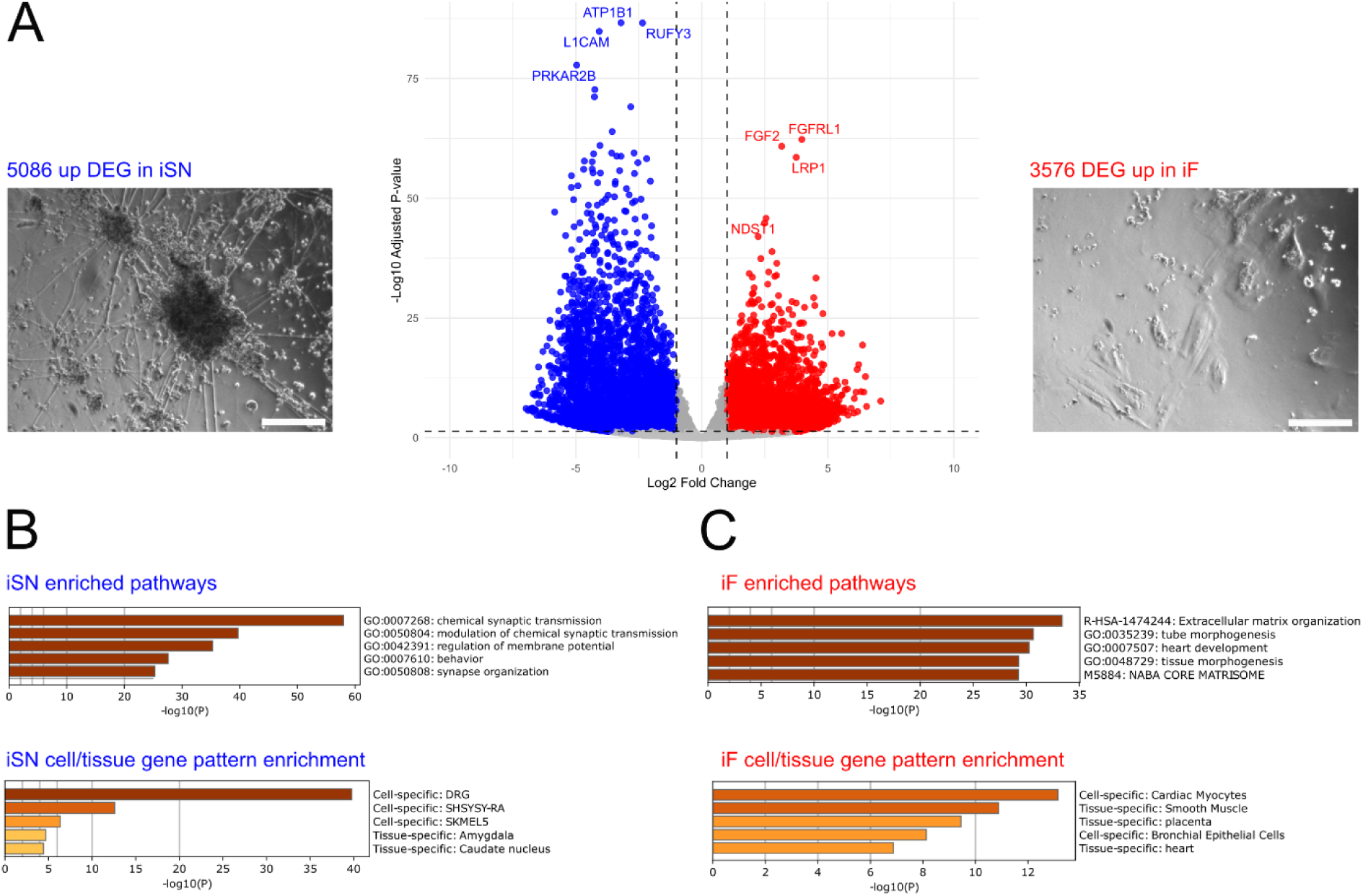
Characterization of CTRL iSN and CTRL ‘feeder-like’ cell transcriptome. A) 5086 genes were upregulated in iSN versus iF, while 3576 genes were upregulated in iF versus iSN. B) ORA and PaGenBase analysis via metascape identified synaptic and membrane potential pathways, emphasizing the DRG origin of iSN. C) For iF, extracellular matrix organization was the most enriched pathway, with best gene set overlap with tissues of cardiac myocytes, smooth muscle, and placenta. Scale bars 200 μm. Comparison between n=4 CTRL iSN differentiations versus n=2 CTRL iF differentiations.

**Supplementary Figure S3.**
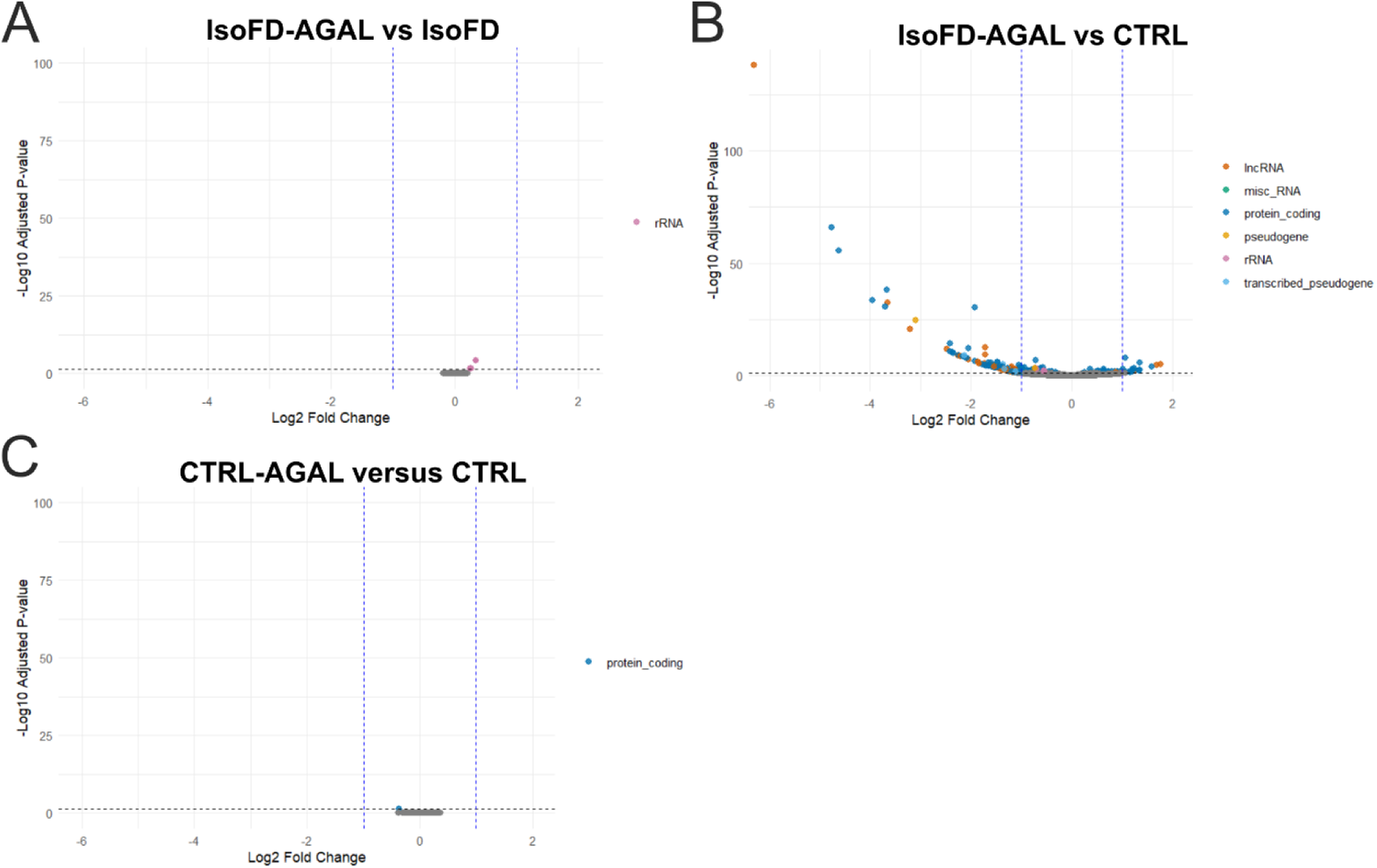
Transcriptional differences related to AGAL treatment. A) IsoFD-AGAL versus IsoFD revealed slight ribosomal RNA upregulation in treated IsoFD. B) IsoFD-AGAL versus CTRL showed persisting dysregulation in treated IsoFD. C) CTRL-AGAL versus CTRL indicated no treatment effect on healthy iSN. Comparison between each n=4 differentiations per cell line and treatment and cutoff at padj<0.05.

**Supplementary Figure S4.**
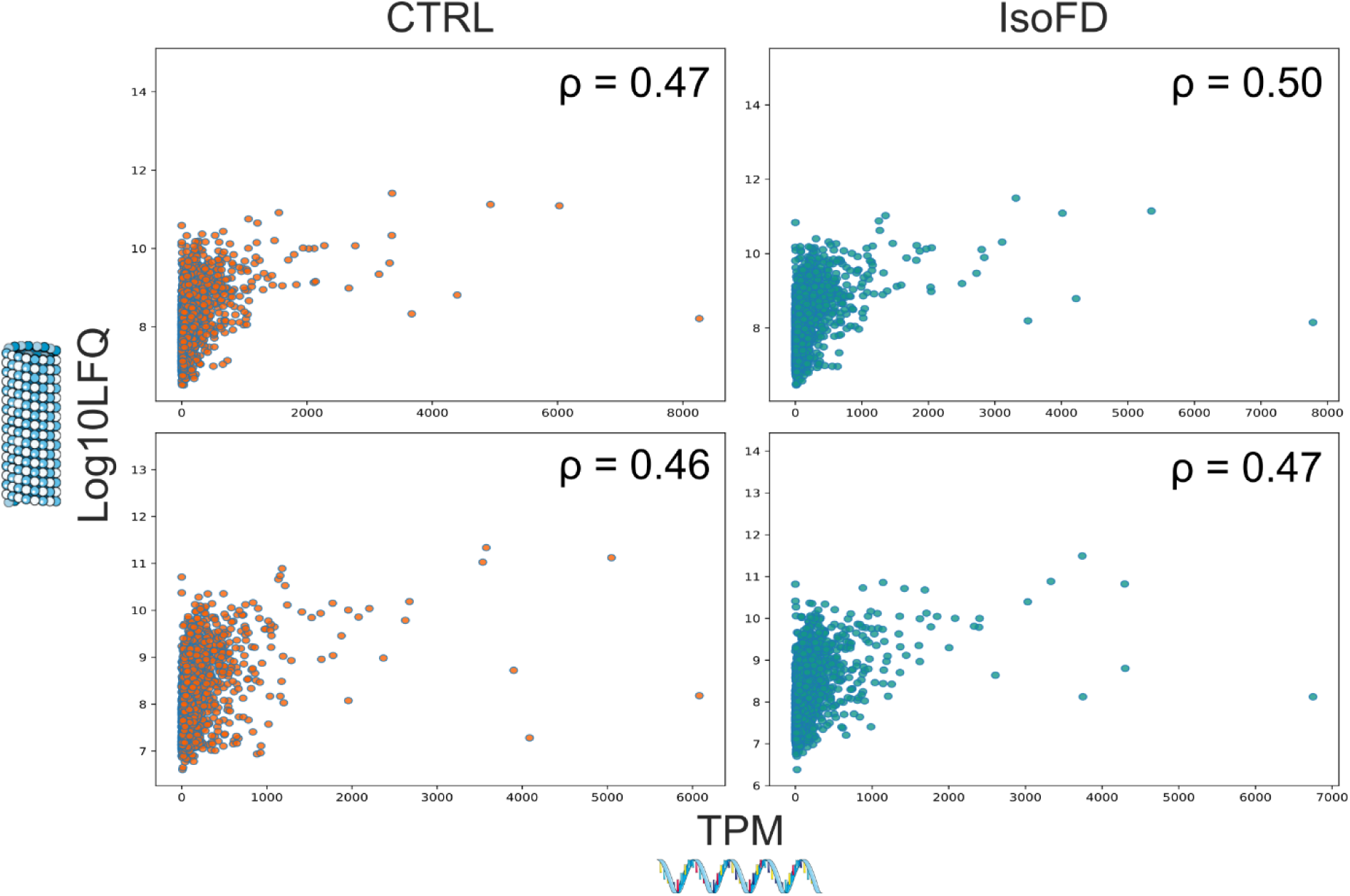
Correlation between RNA transcripts and protein levels for each n=2 differentiations of CTRL and IsoFD iSN.

## Supplementary materials

### Sensory neuron differentiation

During the 10 day differentiation, medium was changed daily. First KO medium (KSR; KO DMEM/F12 + 2 mM GlutaMAX + 15% KO serum replacement + 100 μM 2- mercaptoethanol + 0.1 mM minimum essential medium non-essential amino acids + 100 U/ml pen/strep [all: Thermo Fisher Scientific, Waltham, MA, USA]), containing 100 nM LDN-193189 (STEMCELL Technologies, Vancouver, Canada and 10 μM SB- 431542 (Miltenyi Biotec, Bergisch Gladbach, Germany) (KSR-2i) was used. From day two, medium was supplemented with KSR-2i and 10 μM SU-5402, 10 μM N-[N-(3, 5- difluorophenacetyl)-l-alanyl]-s-phenylglycinet-butyl ester (both: Sigma Aldrich, St.

Louis, MO, USA), and 3 μM CHIR-99021 (Axon Medchem, Groningen, Netherlands) (KSR-2i3i). From day 4, KSR-2i3i medium was replaced by N2 medium [N2; DMEM/F12 GlutaMAX + 1X B-27 Plus Supplement + 1X N-2 Supplement + 100 U/ml pen/strep (all: Thermo Fisher Scientific, Waltham, MA, USA) in 25% increments every second day. On Day 10, cells were splitted with a 1:2 ratio in neuronal maturation medium (N2 medium + 20 ng/ml brain-derived neurotrophic factor (BDNF) + 20 ng/ml glial cell line-derived neurotrophic factor (GDNF) + 20 ng/ml nerve growth factor, beta subunit (NGFb) (all: PeproTech, Rocky Hill, NJ, USA) + 200 ng/ml ascorbic acid (Sigma-Aldrich, St. Louis, MO, USA), and 10 μM floxuridine (FdU, Santa Cruz Biotechnology, Dallas, TX, USA).

### RNA Sequencing

RNA quality was checked using a 5200 Fragment Analyzer with the DNF-471-33 - SS Total RNA 15nt kit (Agilent Technologies). The RQN for all samples was ≥ 7. DNA libraries suitable for sequencing were prepared from 40-70 ng of total RNA with oligo-dT capture beads for poly-A-mRNA enrichment using the TruSeq Stranded mRNA Library Preparation Kit (Illumina) according to manufacturer’s instructions (½ volume). After 16 cycles of PCR amplification, the size distribution of the barcoded DNA libraries was estimated ∼290 bp by electrophoresis on 5200 Fragment Analyzer with the DNF-474-33 - HS NGS Fragment 1-6000bp kit (Agilent Technologies).

Sequencing of pooled libraries, spiked with 1% PhiX control library, was performed at ∼25 million reads/sample in single-end mode with 100 nt read length on the NextSeq 2000 platform (Illumina). Demultiplexed FASTQ files were generated with bcl-convert v4.2.4 (Illumina). To assure high sequence quality, Illumina reads were quality- and adapter-trimmed via cutadapt v2.5 using a cutoff Phred score of 20 in NextSeq mode, and reads without any remaining bases were discarded (command line parameters: --nextseq-trim= 0 -m 1 -a AGATCGGAAGAGCACACGTCTGAACTCCAGTCAC).

Processed reads were subsequently mapped to the human genome (GRCh38.p14 primary assembly and mitochondrion) using STAR v2.7.2b with default parameters based on RefSeq annotation version RS_2023_03 for GRCh38.p14 (Mapping stats of the samples are shown Supplementary Fig. S5). Read counts on exon level summarized for each gene were generated using featureCounts v1.6.4 from the Subread package. Multimapping and multioverlapping reads were counted strand- specific and reversely stranded with a fractional count for each alignment and overlapping feature (command line parameters: -s 2 -t exon -M -O --fraction).

The count output was used to identify differentially expressed genes using DESeq2 version 1.24.0. Read counts were normalized by DESeq2 and fold-change shrinkage was applied by setting the parameter “betaPrior=TRUE. A batch correction was done by including the batch (different differentiations from same cell line with and without AGAL supplementation) in the design formula

(DESeqDataSetFromMatrix(countData=countTable, colData=samples, design=∼batch+condition)).

**Supplementary Figure 5.**
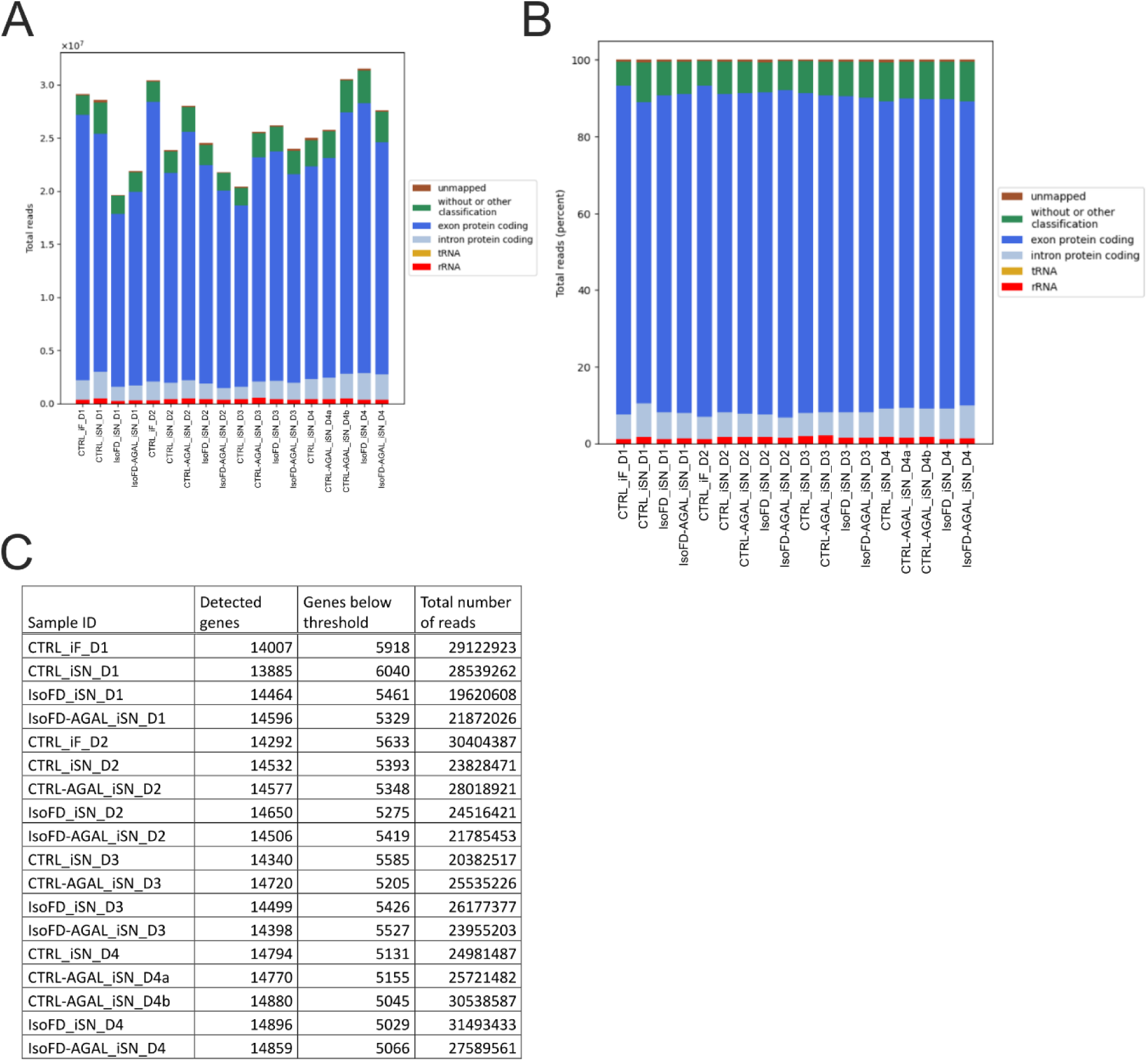
Mapping stats of RNA samples. A) Total read mapping distribution. B) Percentage of read mapping distribution. C) Number of expressed protein coding genes identified (reads mapped >10).

### Proteomics

Protein was precipitated overnight at -20°C with fourfold volume of acetone and afterwards washed three times with acetone at -20°C. Precipitated proteins were dissolved in NuPAGE® LDS sample buffer (Thermo Fisher Scientific, Waltham, MA, USA), reduced with 50 mM DTT at 70°C for ten minutes, and alkylated with 120 mM Iodoacetamide at room temperature for 20 minutes. NuPAGE® Novex® 4-12 % Bis- Tris gels (Thermo Fisher Scientific, Waltham, MA, USA) with MOPS buffer were used for gel electrophoresis according to the manufacturer’s protocol. Gels were washed three times for five minutes with water before staining with Simply Blue™ Safe Stain (Thermo Fisher Scientific, Waltham, MA, USA) for 60 min at RT. Gels were washed with water for one hour and subsequently every gel lane was cut into 15 slices. Gel slices were destained with 30% acetonitrile in 0.1 M NH4HCO3 (pH 8), shrunk with 100% acetonitrile, and dried in a vacuum concentrator (Concentrator 5301, Eppendorf SE, Hamburg, Germany). Gels were digested with 0.1 μg trypsin per gel band overnight at 37°C in 0.1 M NH4HCO3 (pH 8). After removing the supernatant, peptides were extracted from the gel slices with 5% formic acid, and extracted peptides were pooled with the supernatant. Nano liquid chromatography tandem mass spectrometry (Nano-LC-MS/MS) analysis was performed on an Orbitrap Fusion (Thermo Fisher Scientific, Waltham, MA, USA) equipped with a PicoView Ion Source (New Objective, Littleton, MA, USA) and coupled to an EASY-nLC 1000 liquid chromatograph (Thermo Fisher Scientific, Waltham, MA, USA).

Peptides were loaded on a trapping column (2 cm x 150 μm ID, PepSep, Marslev, Denmark) and separated on a capillary column (30 cm x 150 μm ID, PepSep), both packed with 1.9 μm C18 ReproSil (Dr. Maisch HPLC GmbH, Germany) and separated with a 30-minute linear gradient from 3% to 30% acetonitrile and 0.1% formic acid and a flow rate of 500 nl/minute. MS and MS/MS scans were acquired in the Orbitrap analyzer with a resolution of 60,000 for MS scans and 30,000 for MS/MS scans. Higher energy collisional dissociation (HCD) fragmentation with 35% normalized collision energy was applied. A Top Speed data-dependent MS/MS method with a fixed cycle time of three seconds was used. Dynamic exclusion was applied with a repeat count of one and an exclusion duration of 30 seconds; singly charged precursors were excluded from selection. Minimum signal threshold for precursor selection was set to 50,000. Predictive automatic gain control (AGC) was used with an AGC target value of 4×10^5^ for MS scans and 5×10^4^ for MS/MS scans. EASY-IC ion source was used for internal calibration. Raw MS data files were then analyzed with MaxQuant version 1.6.2.2. Database was searched with Andromeda, which is integrated in the utilized version of MaxQuant. The search was performed against the UniProt Human Reference Proteome database (Release 2023_05, UP000005640, 82685 entries). Additionally, the contaminants database integrated in MaxQuant was used. The search was performed with tryptic cleavage specificity and three allowed miscleavages. Protein identification was under control of the false- discovery rate (FDR; <1% FDR on protein and peptide spectrum match [PSM] level). In addition to MaxQuant default settings, the search was performed against following variable modifications: Protein N-terminal acetylation, Gln to pyro-Glu formation (N- term. Gln) and oxidation (Met). Carbamidomethyl (Cys) was set as fixed modification. Further data analysis was performed using R scripts developed in-house. Label-free quantitation (LFQ) intensities were used for protein quantitation.

Proteins with less than two razor/unique peptides were removed. Missing LFQ intensities were imputed with values close to baseline. Data imputation was performed with values from a standard normal distribution with a mean of the 5% quantile of the combined log10-transformed LFQ intensities and a standard deviation of 0.1.

### Fiji pipelines

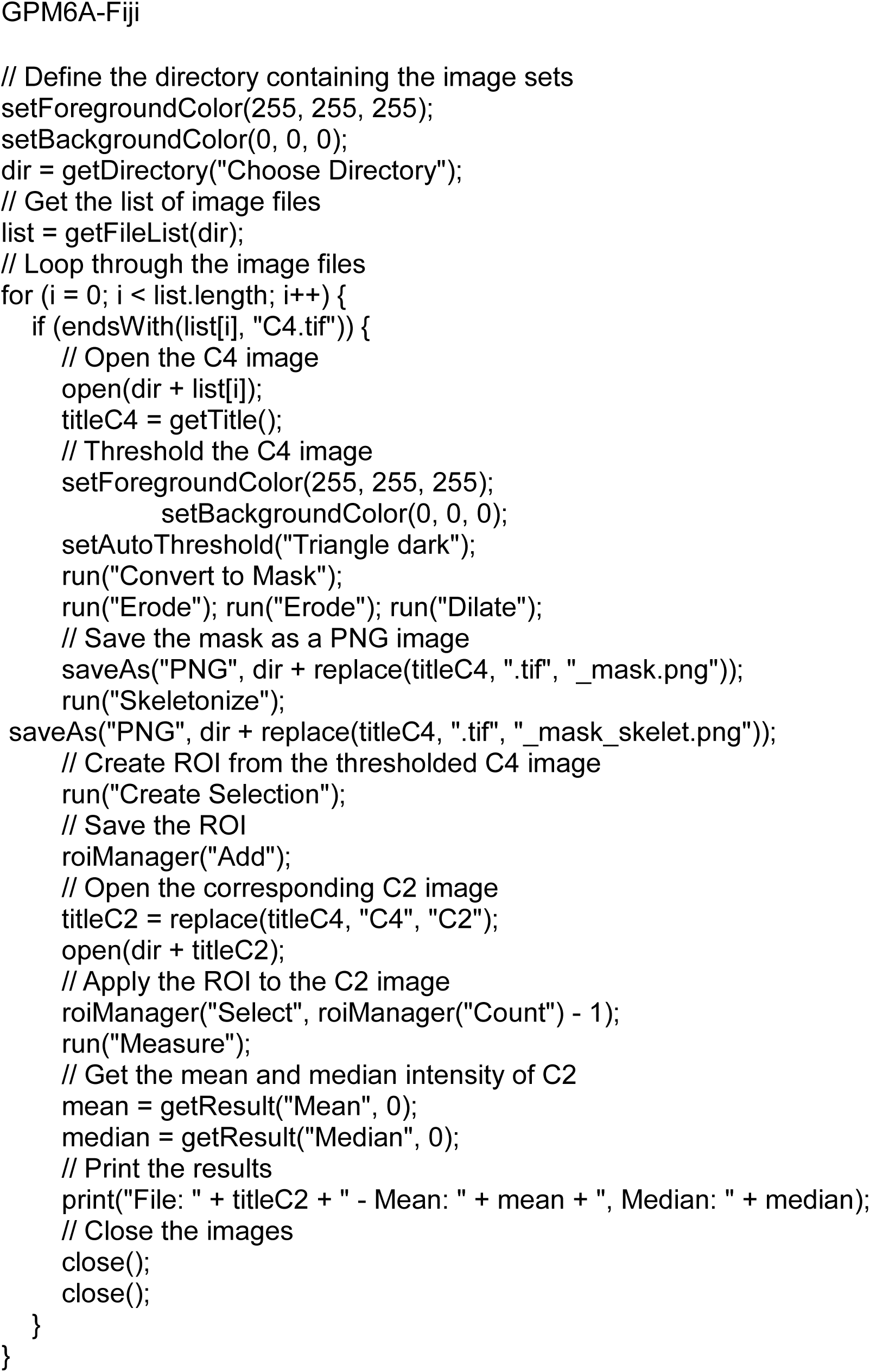
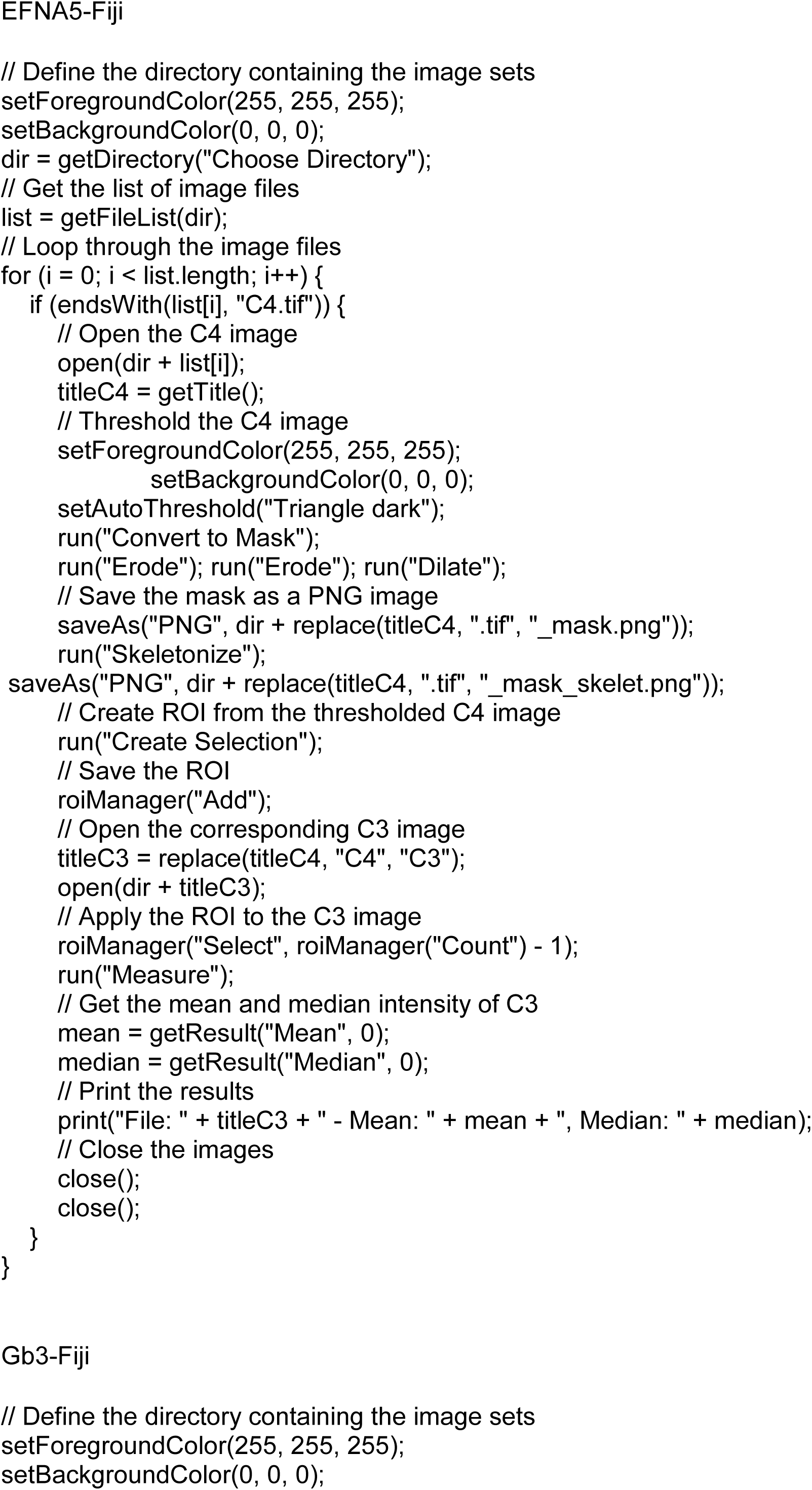
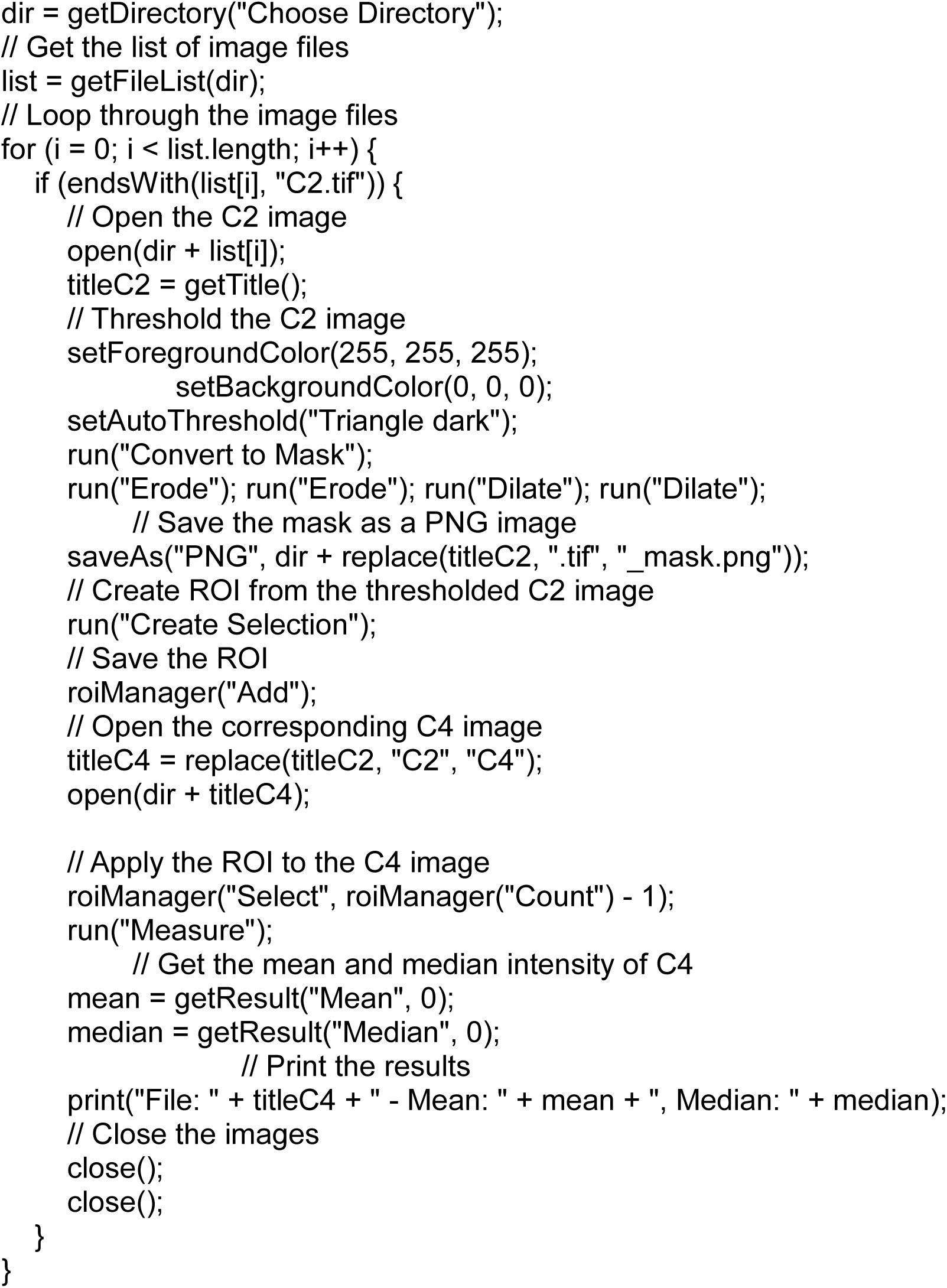

